# Dual forms of aging-related NADPH diaphorase neurodegeneration in the sacral spinal cord of aged non-human primates

**DOI:** 10.1101/527358

**Authors:** Yinhua Li, Zichun Wei, Yunge Jia, Wei Hou, Yu Wang, Shun Yu, Geming Shi, Guanghui Du, Huibing Tan

## Abstract

NADPH diaphorase (NADPH-d) was used to detected neurodegeneration in animal models. In our previous studies, aging-related NADPH-d positive spheroidal bodies (ANB) occurred in the lumbosacral spinal cord of aged rats. Recently, NADPH-d enlarged-diameter neurites named as megaloneurites were detected in the sacral spinal cord of aged dogs. To generalize the occurrence of megaloneurites and/or ANB in non-human primates, the advanced examination of the aging-related phenotypes was undertaken in the monkeys by using the same approach. We discovered two different anomalous NADPH-d positive alterations, which were expressed as ANB and megaloneurites specially distributed in the superficial dorsal horn, dorsal gray commissure, lateral collateral pathway (LCP) and sacral parasympathetic nucleus in the aged monkeys’ sacral spinal cord, compared with the cervical, thoracic and lumbar segments. In the gracile nucleus of aged monkeys, only aging-related spheroidal bodies were observed and no megaloneurites occurred. The dense, abnormal NADPH-d positive megaloneurites, extremely different from regular NADPH-d positive fibers, were prominent in the sacral segments and occurred in extending from Lissauer’s tract through lamina I along the lateral boundary of the dorsal horn to the region of the SPN. Meanwhile, large diameter punctate NADPH-d activity occurred and scattered in the lateral white matter of the LCP and dorsal root entry zone at the same level of NADPH-d abnormality in the gray matters. Those dot-like NADPH-d alterations were examined by horizontal sectioning and indicated ascending or descending oriental fibers. These NADPH-d megaloneurites had the same composition as the punctate NADPH-d alterations and were co-localized with the vasoactive intestinal peptide (VIP) immunoreaction, while the ANB did not coexist with the VIP immunoreaction. Both ANBs and megaloneurites provide consistent evidence that the anomalous neuritic alterations in the aged sacral spinal cord are referred to as a specialized aging marker in the pelvic visceral organs in non-human primates.

## Introduction

The lumbosacral spinal cord different from other segment spinal cord contains autonomic centers regulating the urinary tract, large intestine and sex organs [1–3]. The preganglionic neurons in the intermediolateral nuclei (IML) are very important anatomical structure for the sacral spinal cord [2]. There is also another specialized enlarged area, the dorsal gray commissure (DGC) occurs dorsal of the central canal in the sacral segments. The DGC receives the inputs of pelvic organs and pudendal nerves as well as pelvic nerves through the Lissauer’s tract (LT) and its lateral- and medial-collateral projections [4–9]. Both IML and DGC are part of the reflex pathways of the primary visceral and somatic afferent fibers from the bladder[10–12], external urethral sphincter[11, 13], urethra[14] and the penile nerve[5, 15] as well as colon[16]. In addition, the gracile nuclei receives both ascending projection from the area around the central canal [17] and some primary afferent inputs from pelvic visceral organs[18].

The nicotinamide adenine dinucleotide phosphate-diaphorase (NADPH-d) has been detected histochemically for neuronal nitric oxide synthase (NOS) in the central and peripheral system[19]. Besides labeled perikaryas, NADPH-d positive fibers are distributed more widespread for the powerful enzyme histological visualization. The staining pattern for finest processes of NADPHd-positive cells can be visualized similar to Golgi impregnations and intracellular staining. Both the spinal cord and the gracile nucleus are innervated by NADPH-d neurons and fibers [20]. The NADPH-d positivity distributes in visceral afferent pathway of Lissauer’s tract to the region of the sacral parasympathetic nucleus (SPN) [21–23].

The NADPH-d positivity has been used to study NADPH-d neuritic dystrophy in animal model of Alzheimer’s disease [24]. The significance of NADPH-d for neuronal aging-dependent alterations are elucidated in the sacral spinal cord and gracile nucleus [25–28]. In our previous study, we demonstrate that the age-related NADPH-d positive bodies (ANB) occur specifically in the lumbosacral spinal cord of aged rats[25]. Recently, we also found NADPH-d positive megaloneurits in the sacral spinal cord of aged dogs [26]. Meanwhile, the NADPH-d megaloneurits colocalize with vasoactive intestinal peptide (VIP) [26]. Interestingly, pelvic organs are innervated with VIP [29–31]. On the one hand, aging specialization of NADPH-d alteration implicates aging neurodegeneration in the sacral spinal cord. But on the other hand, ANB in rats is apparently different with specialized megaloneurits in dogs. We wonder if the aging-related NADPH-d alterations occur any other morphological diversity in a higher hierarchy species. Thus, it prompted us to examine whether NADPH-d positive abnormality in the lumbosacral spinal cord of aged non-human primates were consistent with specialized ANBs or megaloneurites.

## Materials and MethodsS

### Animal and tissue preparation

All animal care and treatment in this study were performed in accordance with the guidelines for the national care and use of animals approved by the National Animal Research Authority (P.R. China).Young (less than 5-year-old, n=10) and aged (more than 15~29-year-old, n=12) cynomolgus monkeys of both sexes were used in our experiments. All animals were housed under a 12:12-h light/dark cycle in the animal care facilities. The facilities are certified by the Council on Accreditation of the Association for Assessment and Accreditation of Laboratory Animals Care (International) and or China National Accreditation Service for Conformity Assessment. The ambient temperature was maintained at 24±2 °C and relative humidity at 65±4%. Reverse osmosis water was available *ad libitum*. Normal food, fresh fruit and vegetables were supplied twice daily. Animals were purchased from Beijing Xierxin Biological Resources Research Institute (Permit Number: 11805300010425, 11805300010433, Beijing, China), Wincon TheraCells Biotechnologies Co., Ltd. (Permit Number: WD-0312010, Naning, China) and Hainan Jingang Biotech Co., Ltd (Permit Number: 46001200000114, Hainan, China). All experimental procedures were approved by the Institutional Ethics Committee in Animal Care and Use of the Jinzhou Medical University (approval ID: 2014JCATBH0E-NNSF-0004).

The animals were anesthetized with sodium pentobarbital (50 mg/kg i.p.) and perfused transcardially with saline followed by 4% paraformaldehyde in a 0.1M phosphate buffer saline (PBS, pH 7.4). Following perfusion fixation, the spinal cords and brains were obtained and placed in 30% sucrose for 48~72 hrs. The spinal cords of the all segments and gracile nucleus in the medulla oblongata were cut transversely of 40μm sections on a cryostat. To visualize the rostrocaudal orientation of the NADPH-d positivity, horizontal sections (40μm) of the spinal cords and medulla oblongata of the aged monkeys were also performed.

### NADPH diaphorase histochemistry

Free floating sections were used for staining [25]. Most of the tissue sections from the young and old monkeys were stained by NADPH-d histochemistry, with incubation in 0.1 M PBS (PH 7.4), 0.3% Triton X-100 containing 1.0 mM reduced-NADPH (Sigma, St. Louis, MO, USA) and 0.2 mM nitro blue tetrazolium (NBT, Sigma), at 37°C for 2 to 3 h. The reaction was stopped by washing the sections with 0.1 M PBS.

### Double Immunofluorescence staining

Subsequently, some sections were performed by double-staining with NADPH-d histochemistry and CGRP, VIP or NPY immunofluorescence, respectively. The sections were collected in 24-well plates containing 0.1 M PBS and washed three times. Then sections were incubated for 1 h at room temperature in blocking solution (0.1 M PBS) at pH 7.4 with 1% BSA and processed for free-floating immunofluorescence using primary polyclonal antibodies that label calcitonin gene-related peptide (CGRP, mouse; 1:100, Sigma, USA), vasoactive intestinal peptide (VIP, rabbit, 1:1000 Sigma, USA), neuropeptide Y (NPY, rabbit; 1:5000, Sigma, USA). The sections were incubated with the primary antibody in PBS containing 1% BSA for 24 h at 4 °C, respectively. For a negative control, a few of sections were incubated without primary antibody in each immunofluorescence testing. The sections were then washed several times with PBS. Fluorescent-conjugated secondary antibodies (Goat anti-Rabbit IgG (H+L), Alexa Fluor 594 [1:800, Life] and Goat anti-Mouse IgG, Alexa Fluor 488 [1:800, Life]), were applied to the sections for 1 h at 37 °C in the dark. Finally, after several washes with PBS, the sections were counterstained with DAPI for 10 min. The sections were mounted onto slides and coverslipped. For controls of immunofluorescence staining omitted the primary antibody, no specified immunostaining was detected in the control experiments.

### Image acquisition and Figure edition

Images were captured with a DP80 camera in an Olympus BX53 microscope (Olympus, Japan). Figures were edited with CorelDraw (Ottawa, Canada).

## Result

### NADPH diaphorase activity in the sacral spinal cord of aged monkeys

According our previous experiments [25, 26], the sacral segments and gracile nucleus were focused, especially the dorsal root entry zone (DREZ), lateral collateral pathway (LCP), SPN and DGC where the NADPH-d positive megaloneurites and ANB distributed in aged monkeys. The NADPH-d staining revealed a moderately dense fiber network of dendrites and axon terminals as well as moderately stained, small- and medium-sized neurons in the dorsal horn, DREZ, LCP and DGC of sacral spinal cord of young monkeys (Fig. 1A, B). When compared with young monkeys (Fig. 1A, B), the NADPH-d positive megaloneurites were observed in the DGC and the LCP of LT (Fig. 1C, D) and the ANB were mainly distributed in the SPN and superficial dorsal horn of the sacral spinal cord of aged monkeys (black arrowheads in Fig. 1C). The general location and selective segmental distribution of the NADPH-d positive megaloneurites and the ANB were related to the central projection of the primary pelvic visceral sensation, mostly located dorsal of the spinal cord. Further segmental examination was to demonstrate that the megaloneurites occurred in a specific regional localization. In transverse sections of caudal spinal segments, NADPH-d stained fiber network of dendrites and axon terminals were noted in the superficial dorsal horn (laminae I, II), the LCP of LT, within the dorsal lateral funiculus. As mentioned above, the most prominent fiber staining in the sacral segments of aged monkeys was in LT along the lateral edge of the dorsal horn to the region of the SPN (Fig. 1A, C). Megaloneurites were selectively detected in the sacral spinal cord but not in lumbar or in thoracic and cervical segments (Fig. 1I-K).

**Fig. 1.**
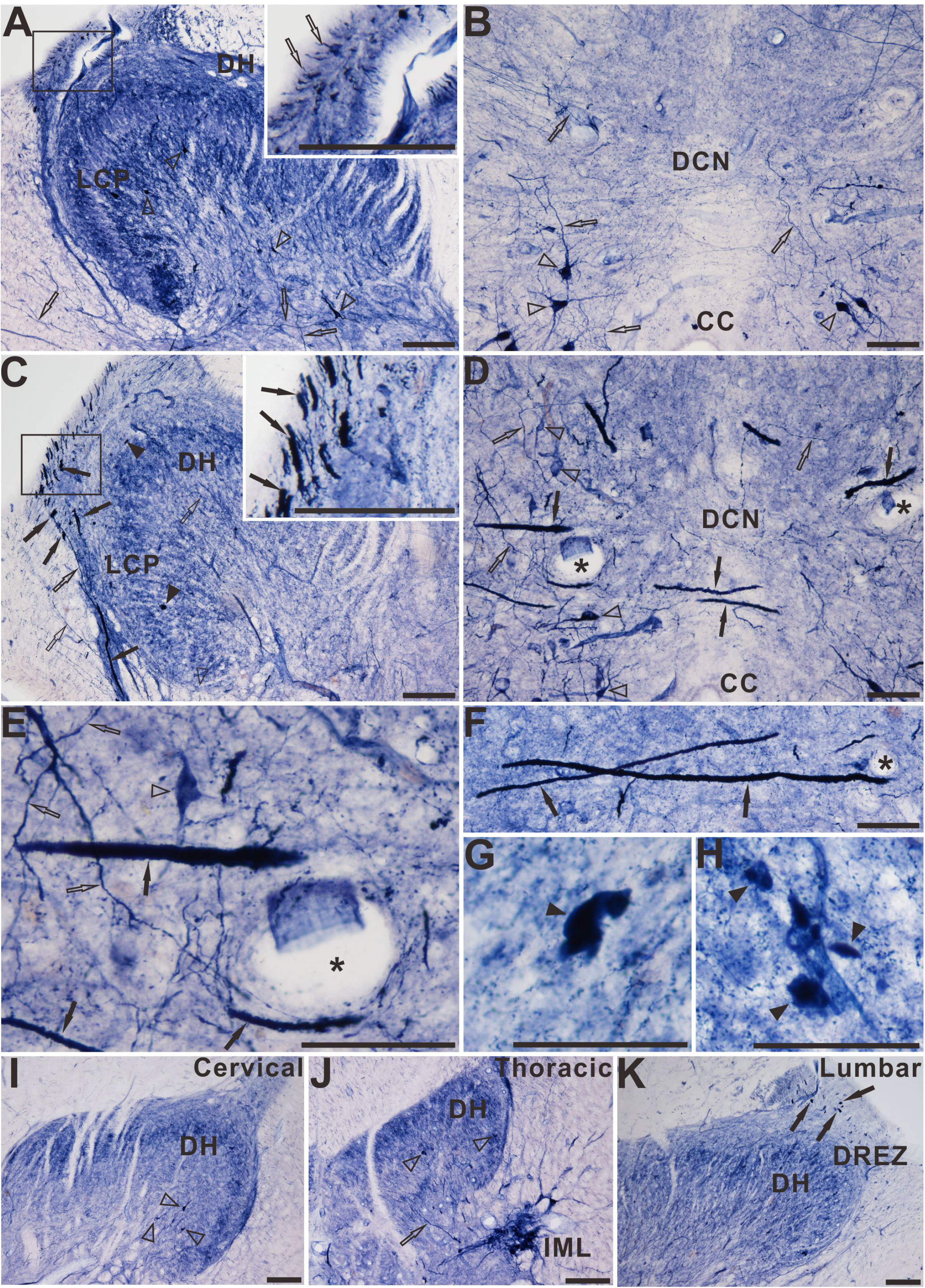
Microphotographs of NADPH-d positive reactivity in young (**A, B**) and aged (**C, D**) monkeys at the sacral spinal cord. All of the transverse sections taken at the same levels. The illustration represents the magnification of the corresponding rectangular frame. The NADPH-d positive megaloneurites and ANBs in the sacral spinal cord of aged monkeys are completely different from the surrounding normal fibers and neurons (**E-H**).The NADPH-d positive activity in the dorsal horn of cervical (**I**), thoracic (**J**) and lumbar (**K**) segment of aged monkeys. The asterisks indicate vascular structures. Open arrowheads: NADPH-d positive neurons, black arrowheads: ANBs, open arrows: normal NADPH-d positive neurites, black arrows: megaloneurites. Scale bar in **A, C, I-K**=100μm, in **B, D, E-H**= 50μm

### NADPH-d activity in the white matter and the DREZ

In the sacral spinal cord of aged monkeys, enlarged massive NADPH-d activity (Fig. 2A) in transverse sections was detected in the white matter compared with young monkeys (Fig. 2B). It was confirmed similar to megaloneurites in transverse section, which may be possibly in close association with NADPH-d positive megaloneurites penetrating deeply into the white matter. Considerably higher punctate NADPH-d activity was detected in the lateral portion of the LCP, and the abnormal alterations were extremely different from normal nerve cells (Fig. 2A). Horizontal sections of the sacral segment (Fig. 2C) indicated that the punctate NADPH-d alterations were longitudinally-arranged fibrous extending rostrocaudally and were greatly different from normal fiber bundles and vascular structures in white matter (Fig. 2C). In addition, numerous NADPH-d fibers and varicosities were evident in LT and in some instances, these fibers could be traced along the whole length of the section (Fig. 2D). We still termed the aged and segmental associated alterations in the white matter as megaloneurites.

**Fig. 2.**
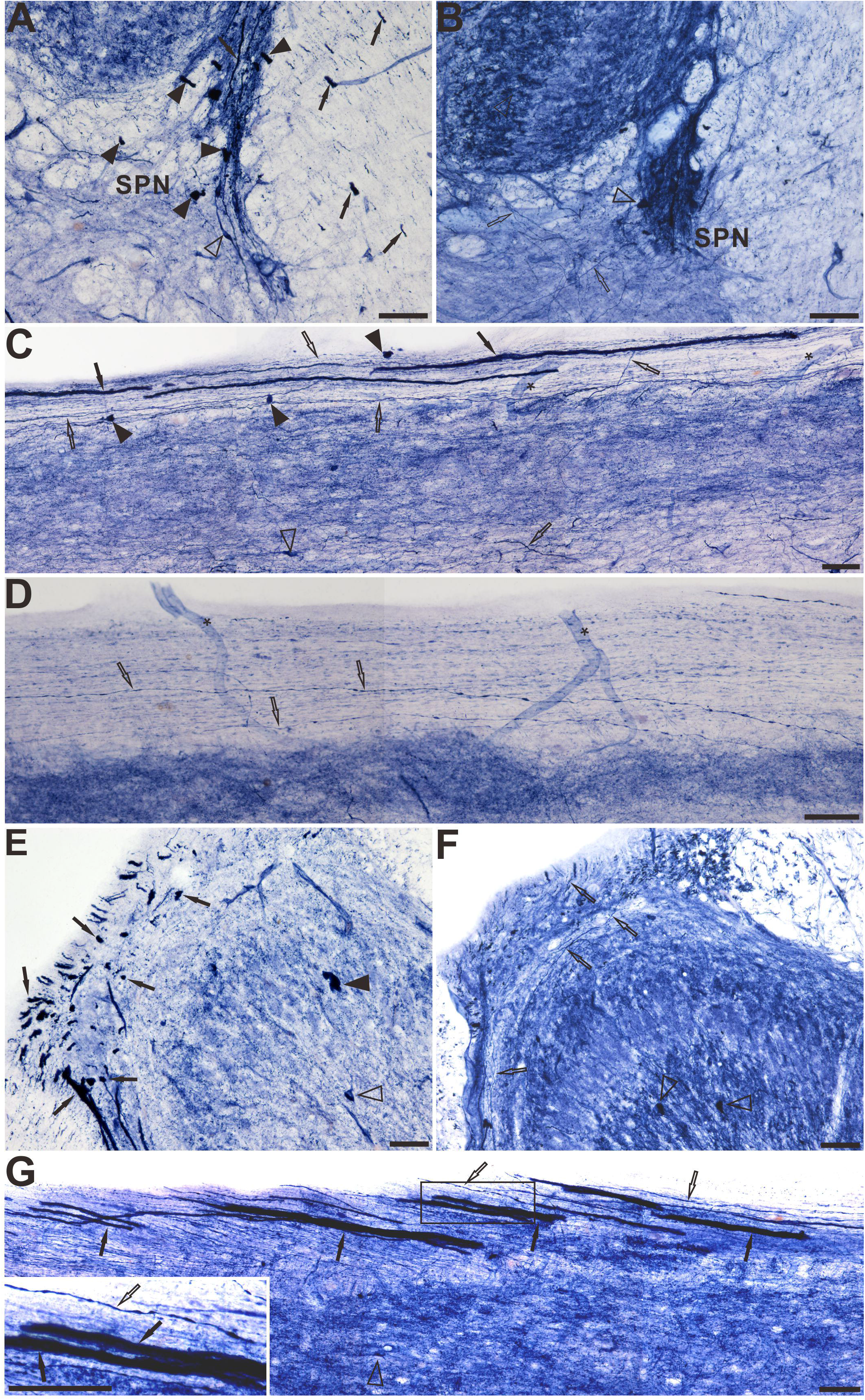
The NADPH-d positive alterations in the white matter and DREZ of the sacral spinal cord of monkeys. **A** indicates that the location and morphology of the NADPH-d positive megaloneurites and ANBs in the white matter and LCP in sacral segment of aged monkeys and **B** represents the NADPH-d positive activity of young monkeys. In the horizontal section, NADPH-d staining fibers in the white matter of the lateral fasciculus of the aged monkeys (C) and young monkeys (**D**). The distribution of NADPH-d positivity in the DREZ of the sacral spinal cord of aged monkeys (**E**) and young monkeys (**F**). Horizontal section of aged monkeys (**G**) at the sacral segment level showing aberrant NADPH-d positive megaloneurites in the DREZ, transitionally continuous with LT. The illustration represents the magnification of the corresponding rectangular frame. Asterisks indicate vascular structures. Open arrowheads: NADPH-d positive neurons, black arrowheads: ANBs, open arrows: normal NADPH-d positive neurites, black arrows: megaloneurites. Scale bar=50μm

Transverse sections in the DREZ at sacral segment of aged monkeys (Fig. 2E), showing numerous dot-like intensely NADPH-d activities accompanied with the NADPH-d megaloneurites in the LCP. These punctate abnormalities occurred specifically in the caudal spinal cord of aged and not appeared in young animals (Fig. 2F). In horizontal sections, these punctate NADPH-d alterations had the same morphology as NADPH-d positive megaloneurites (Fig. 2G). The properties of the punctate anomalous structures in the horizontal section were that afferent fibers from the dorsal rootlets undergo some pathological changes in the DREZ, forming NADPH-d megaloneurites that continue with LT (Fig. 2G). The enlarged neurites were not found in the distal rootlets and DRGs in the aged monkeys. Compare to the other segments of the spinal cords, the megaloneurites mainly occurred in the DREZ of sacral segment, occasionally in the lumbar and rostral coccygeal in aged monkeys.

### Double staining with three neuropeptides

The double-staining of NADPH-d histochemistry combined with CGRP, Neuropeptide Y and VIP immunofluorescence were used to identify the megaloneurite properties respectively (Fig 3). No structures corresponding to NADPH-d positive megaloneurites were detected by CGRP, Neuropeptide Y immunofluorescence (Fig. 3A-F). For VIP immunoreactivity, VIP and NADPH-d mageloneurites positively localized in DGC (white arrowheads in Fig. 3G-I). The ependymal cells of central canal were detected with VIP immunoreactivity (Fig. 3 H, I). In addition, double-staining experiments showed that the punctate NADPH-d alterations in the white matter and DREZ were marked by VIP immunofluorescence and not marked by CGRP or Neuropeptide Y immunofluorescence (fig. 4). The double-staining of NADPH-d histochemistry combined with VIP immunofluorescence displayed that the ANBs distributed in the superficial dorsal horn and SPN were not colocalized with the VIP immunofluorescence, indicating that the composition of the ANB was different from the NADPH-d positive megalonerites and they were two different degenerative structures appearing in the lumbosacral segments.

**Fig. 3.**
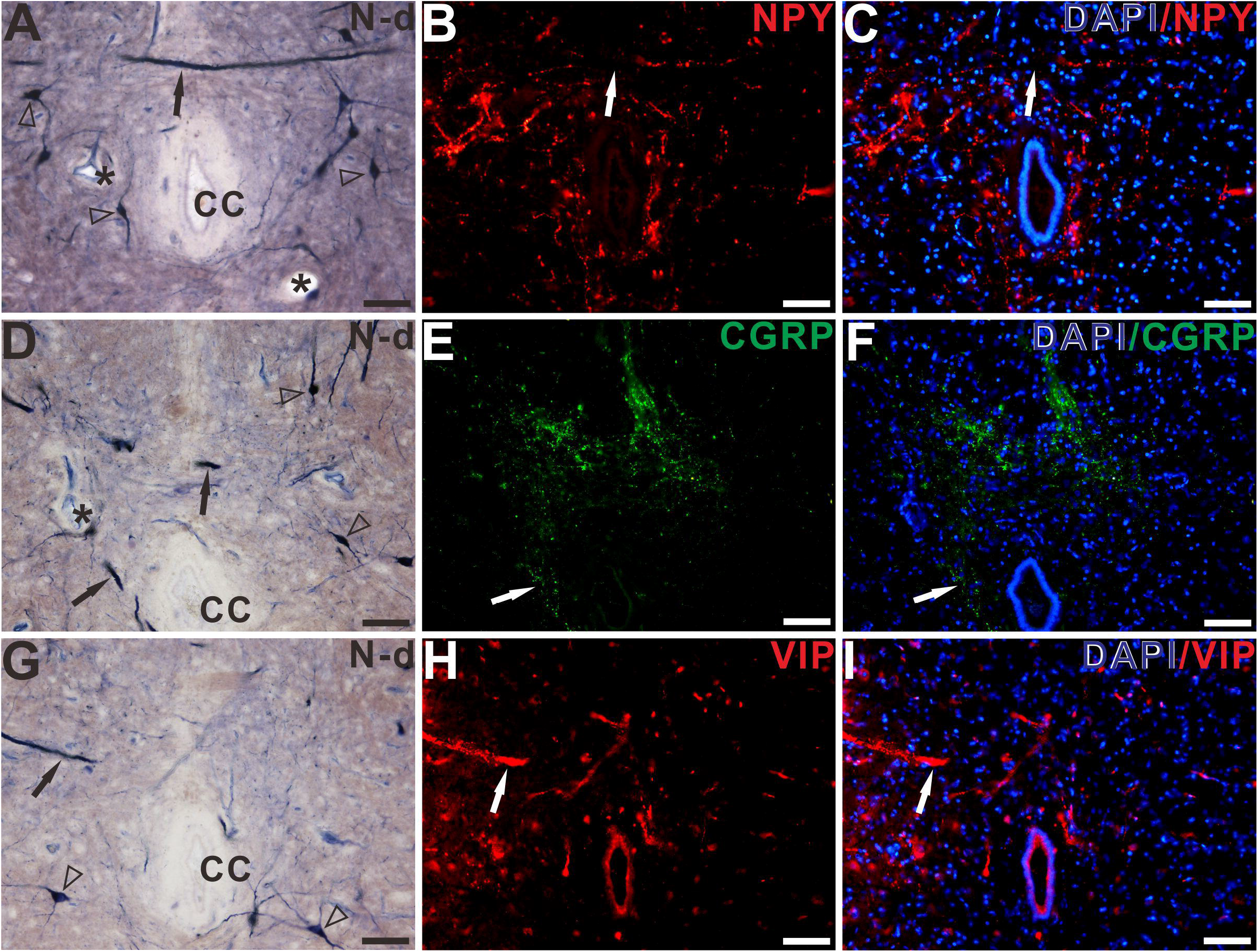
The double-staining of NADPH-d histochemistry combined with Neuropeptide Y (**A-C**), CGRP (**D-F**) and VIP (**G-I**) immunofluorescence in the DCN of sacral segments of aged monkeys. VIP is colocalized with megaloneurites (**G-I**). While Nueropeptide Y and CGRP failed to label the megaloneurites (**A-F**). Asterisks indicate vascular structures. Black/white arrows: megaloneurites, open arrowheads: normal NADPH-d positive neurons. Scale bar=50μm

**Fig. 4.**
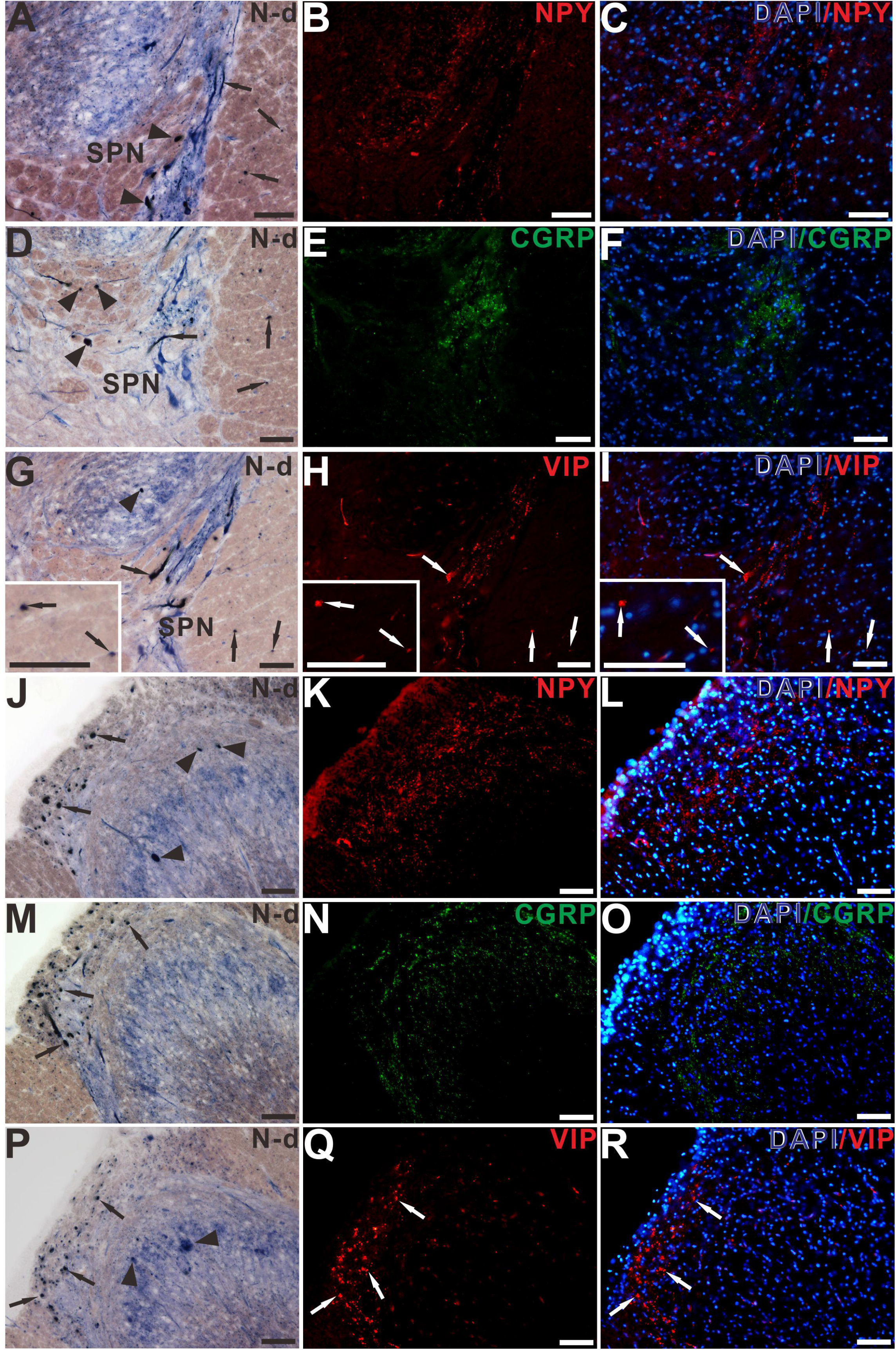
The double-staining of NADPH-d histochemistry combined with Neuropeptide Y, CGRP and VIP immunofluorescence in the white matter (**A-I**) and DREZ (**J-R**) of sacral segments of aged monkeys. The punctate NADPH-d alterations in the white matter (**G-I**) and DREZ (**P-R**) were marked by VIP immunofluorescence and not marked by CGRP or Neuropeptide Y immunofluorescence. The ANBs (black arrows) distributed in the superficial dorsal horn and SPN were not colocalized with the VIP immunofluorescence. Black arrowheads: ANBs, black/white arrows: megaloneurites. Scale bar=50μm

### NADPH-d activity in the caudal medulla

In the caudal medulla of aged monkeys, primarily in the gracile nucleus and cuneate nucleus (Fig. 5A-D), a large number of spheroidal ANBs were detected compared with young monkeys (Fig. 5E-H). These results coincide with those obtained from previous studies in the gracile nucleus of aged rats [27, 32]. ANBs in the gracile nucleus and cuneate nucleus failed for VIP immunoreactivity (data not showed here).

**Fig. 5.**
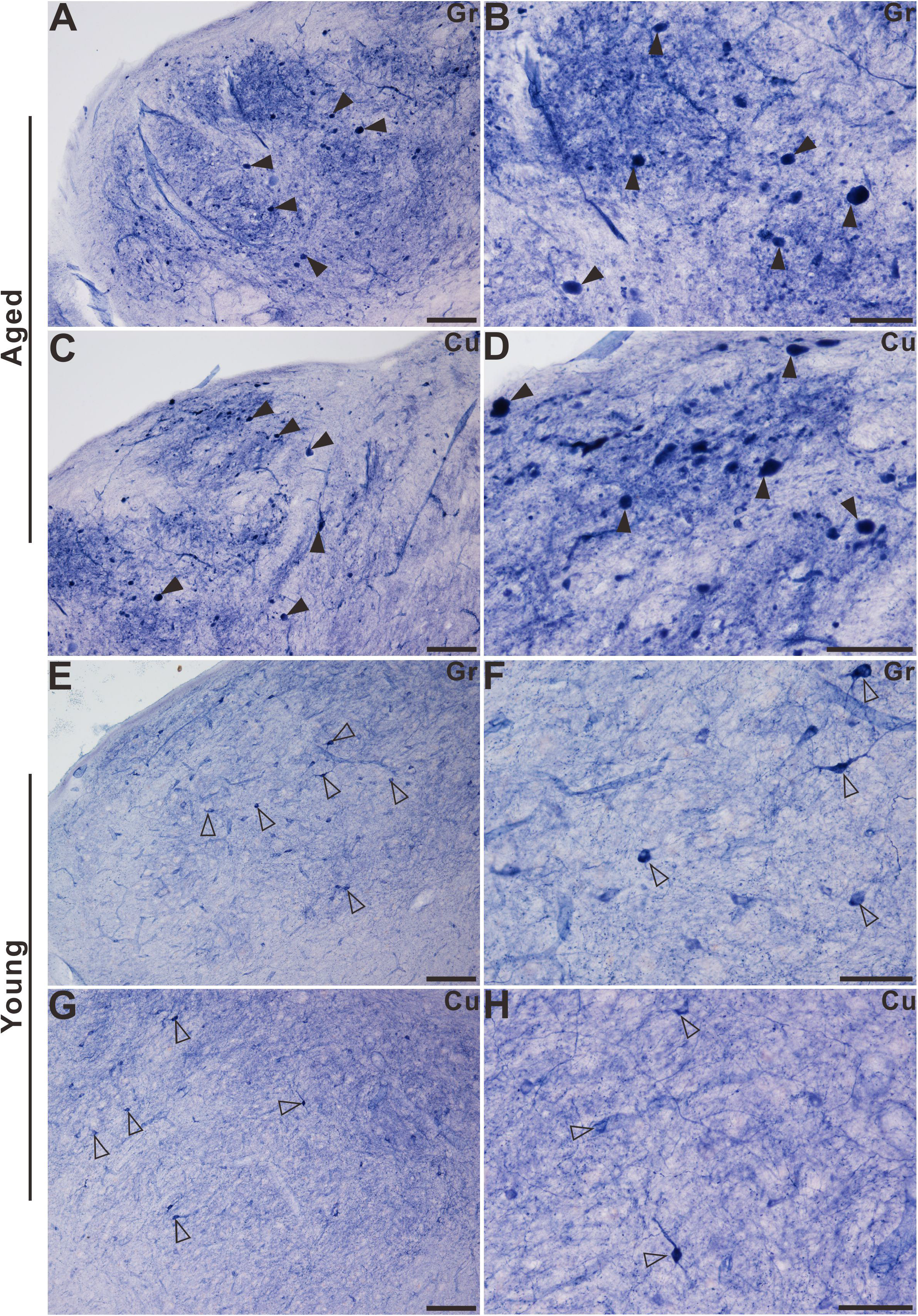
Microphotographs of NADPH-d positive reactivity in the gracile nucleus and cuneate nucleus of aged (**A-D**) and young (**E-H**) monkeys. Note abnormal dot-like NADPH-d positive bodies (black arrowheads) in the gracile nucleus and cuneate nucleus of aged monkeys (**A-D**). The regular NADPH-d positive neurons were observed in the gracile nucleus and cuneate nucleus of young monkeys (**E-H**). Open arrowheads: NADPH-d positive neurons, black arrowheads: ANB. Scale bar in **A, C, E, G**=100μm, in **B, D, F, H**=50μm.

## Discussion

Many studies have proven that NADPH-d reactivity occurs extensively in the spinal cord neurons and sensory pathway [23, 33–40]. Our previous research showed that ANBs occurred in the lumbosacral spinal cord of aged rats [25]. Our recent research also demonstrated that a novel NADPH-d positive megaloneurites in the dorsal horn and DGC of the sacral spinal cord of aged dogs was found to be extremely different from regular NADPH-d positive fibers, especially in the LCP of sacral segments [26]. Although the normal and some experimental NADPH-d positivity were studied in the monkey spinal cord [41], little report was available on the aging related changes.

The major discovery of this investigation was the dual features of ANBs and megaloneurites detected in the lumbosacral spinal cord and ANBs in gracile nucleus of the aged monkeys, and demonstrated their basic morphological property, segmental and laminar distribution. The megaloneurites were different from the previously reported meganeurites which occurred proximal to axonal initial segments of the somas in younger organisms [42]. The anatomical locations are also strikingly different between the two structures. The megaloneurites were assumed as swelling neuronal processes, which were enlarged-diameter formations much longer than meganeurites [42]. Normal NADPH-d positive fibers stained with clear punctate and considerable varicosities. Megaloneurites were intensively stained with less varicosities, and clearly traceable for substantial distances in sacral segments, especially in the horizontal sections.

One of the differences between dogs and monkeys was the lack of NADPH-d positive cells in the region of the SPN of the dogs. In the monkeys, a large percentage of preganglionic neurons was revealed in the region of the SPN, while a few scattered NADPH-d cells were detected in that of the dogs [26]. The NADPH-d positive megaloneurites with morphology and organization were first discovered in the sacral spinal cord in aged dogs [26]. Our study revealed a special occurrence of the aged-related NADPH-d positive megaloneurites and spheroidal ANBs in dorsal part of the lumbosacral spinal cord of aged monkeys. Similar to the experimental results in the sacral spinal cord of aged dogs [26], the dense hypertrophic megaloneurites extending from DREZ through LT along the lateral edge of the dorsal horn to the SPN and DGC in the sacral spinal cord of aged monkeys. In this part of the fibers might play a role in visceral reflex pathways [9, 43], because the sacral DGC and the LCP receives terminations from the somatic and visceral afferents [6, 7, 9, 44]. Functionally, the sacral spinal cord was known to be associated with bowel bladder, and sexual dysfunction [45–47]. The distribution of the megaloneurites overlapped both the efferent and afferent pathways of the autonomic system, which regulates the pelvic organs. Previous tracing experiments in the rat have also demonstrated that a large percentage of afferent neurons projecting to pelvic visceral organs [12, 36] and specifically to the bladder and urethra [12, 21] exhibit NADPH-d activity.

The megaloneurites were confirmedly detected in both coronal and horizontal sections as longitudinally extending fiber bundles mostly in the lateral white matter in the caudal sacral spinal cord. On the basis of these profiles, we suggest that they are a part of the neurites in the lumbosacral dorsal spinal cord. As for their origin, the aging-related megaloneurites in aged dogs were clearly distinguished from endothelium, as the morphology of typical ones was significantly different from the vascular structures found under the light microscopy [25]. Double-staining experiment demonstrated that VIP and NADPH-d megaloneurites positively localized in the DREZ, lateral white matter of the LCP, DGC and LCP. We postulate that the degeneration of NADPH-d fibers happened in the sacral aging condition.

The megaloneurites evidently occurred DREZ by horizontal sections of aged monkeys. By analysis of the horizontal sections, it is consistent that diameter of neurites are gradually enlarged from distal to proximal portions through the DREZ [26], which is a transitional location between peripheral nervous system and CNS in the spinal cord and brain stem [48].

All these of segmental selective deteriorations demonstrated that the sacral segments exited relevant adverse conditions more vulnerable to aging degenerations. The pelvic visceral sensory information from the urogenital organs is routed to the sacral spinal cord through centrally-projecting axons that pass into the CNS via the DREZ and continue with LT. DREZ plays important role in neurodegeneration [49] and axon routing during development [50].

Aging neuropathological studies of NADPH-d positivity could be used to reveal the neuronal terminal-pathy of aged conditions [27] and neurodegenerative animal models [24]. In our unpublished study, the aging related NADPH-d alteration was detected in the gracile nucleus in aged rats [32]. We also detected spherical aberrant NADPH-d positive alterations in the gracile nucleus of aged monkeys. We presume that these spherical alterations were identical to the ANB in aged rats, which is similarly derived from the aging alterations in the lumbosacral spinal cord [25, 51]. The central-projecting axons of the first-order sensory neurons in DRG could configure bifurcation in the spinal cord. The bifurcating branches could terminal in the corresponding spinal segments and ascend to dorsal column nuclei respectively [52]. Beside somatic sensory inputs, dorsal column nuclei also receive visceral sensory inputs [53]. The spheroidal NADPH-d neurodegenerations in aged rats and monkeys were postulated as dying back in the lumbosacral spinal cord and gracile nucleus [54, 55]. We hypothesize that the megaloneurites may be relevant to the mechanism of dysfunction urogenental organs in aged human.

## Conclusion

In summary, the major new finding revealed by this study is that the NADPH-d positive megaloneurites and ANB occurred in the aged lumbosacral spinal cord, but not in young monkeys. These results confirms our discovery in aged rats and dogs, respectively. In addition, NADPH-d megaloneurites could be also labeled with VIP immunofluorescence, while ANB were scarcely labeled, indicating that megaloneurites and ANB were two different neurodegenerations. Our results suggested that the megaloneurites and ANB are considered as specific aging markers, associated progressive aging deterioration and malformed structures.

## Acknowledgments

This work was supported by grants from National Natural Science Foundation of China (81471286), Undergraduate Training Programs for Innovation and Entrepreneurship of Liaoning (201410160007) and Research Start-Up Grant for New Science Faculty of Jinzhou Medical University (173514017). Authors are grateful Dr. Xintian Hu to provide tissue for our experiment and Ms. Wei Wu, Huan Liu, Fengming Gao, Xuecao Tong, Ximeng Xu, Mr. Zhongyuan Xia and Chenxu Rao for technique assistant.

## Author Contributions

YL and ZW conceived and performed the experiments as well as analyzed data. YJ, WH, YW, SY, SY, GS, HT assisted YL and ZW in the experimentation. GD and HT provided important experimental guidance. HT, SY and GD discussed the results. YL and HT wrote the manuscript. HT supervised the project and coordinated the study.

